# Astrocyte store-operated calcium entry is required for centrally mediated neuropathic pain

**DOI:** 10.1101/2023.06.08.544231

**Authors:** Mariya A. Prokhorenko, Jeremy T. Smyth

**Affiliations:** Neuroscience Graduate Program; Department of Anatomy, Physiology, and Genetics, F. Edward Hébert School of Medicine, Uniformed Services University of the Health Sciences, Bethesda, MD

**Author notes:** Address for Correspondence: Jeremy T. Smyth Department of Anatomy, Physiology, and Genetics Uniformed Services University 4301 Jones Bridge Rd Bethesda, MD 20814.

## Abstract

Central sensitization is a critical step in chronic neuropathic pain formation following acute nerve injury. Central sensitization is defined by nociceptive and somatosensory circuitry changes in the spinal cord leading to dysfunction of antinociceptive gamma-aminobutyric acid (GABA)ergic cells (Li et al., 2019), amplification of ascending nociceptive signals, and hypersensitivity (Woolf, 2011). Astrocytes are key mediators of the neurocircuitry changes that underlie central sensitization and neuropathic pain, and astrocytes respond to and regulate neuronal function through complex Ca^2+^ signaling mechanisms. Clear definition of the astrocyte Ca^2+^ signaling mechanisms involved in central sensitization may lead to new therapeutic targets for treatment of chronic neuropathic pain, as well as enhance our understanding of the complex central nervous system (CNS) adaptions that occur following nerve injury. Ca^2+^ release from astrocyte endoplasmic reticulum (ER) Ca^2+^ stores via the inositol 1,4,5-trisphosphate receptor (IP^3^R) is required for centrally mediated neuropathic pain (Kim et al, 2016); however recent evidence suggests the involvement of additional astrocyte Ca^2+^ signaling mechanisms. We therefore investigated the role of astrocyte store-operated Ca^2+^ entry (SOCE), which mediates Ca^2+^ influx in response to ER Ca^2+^ store depletion. Using an adult *Drosophila melanogaster* model of central sensitization based on thermal allodynia in response to leg amputation nerve injury (Khuong et al., 2019), we show that astrocytes exhibit SOCE-dependent Ca^2+^ signaling events three to four days following nerve injury. Astrocyte-specific suppression of Stim and Orai, the key mediators of SOCE Ca^2+^ influx, completely inhibited the development of thermal allodynia seven days following injury, and also inhibited the loss of ventral nerve cord (VNC) GABAergic neurons that is required for central sensitization in flies. We lastly show that constitutive SOCE in astrocytes results in thermal allodynia even in the absence of nerve injury. Our results collectively demonstrate that astrocyte SOCE is necessary and sufficient for central sensitization and development of hypersensitivity in *Drosophila*, adding key new understanding to the astrocyte Ca^2+^ signaling mechanisms involved in chronic pain.

## Results

### Central sensitization following leg amputation injury in adult flies

We developed methods for modeling central sensitization and neuropathic pain in adult *Drosophila* based on Khuong et al (Khuong *et al*., 2019) and adapted these methods for the analysis of astrocyte Ca^2+^ signaling mechanisms. These methods are based on the development of thermal allodynia, a form of hypersensitivity or chronic pain in which a normally non-painful temperature is perceived as painful. Adult flies exhibit escape behaviors in the form of jumps when their legs are in contact with surface temperatures that they perceive as noxious or painful. As shown in Figure 1A-C and Supplemental Figure S1, healthy uninjured animals exhibit relatively few jumps on surface temperatures from 24° C to 38° C, whereas repeated jumping behavior is exhibited by animals on a 42° C surface. This indicates that in healthy flies 42° C is perceived as painful while 38° C is below the threshold for pain sensation. Jumping at 42° C was completely inhibited in animals with *ppk-GAL4*-driven, sensory neuron-specific suppression of the thermal-sensitive TrpA1 ion channel (Supplemental Figure S1), further verifying that jumping is a response to thermal nociception. We then introduced nerve injury in the form of a single middle leg amputation as illustrated in Figure 1A and quantified jumping behavior at the subthreshold 38° C temperature to assess the development of central sensitization and thermal allodynia. As shown in Figure 1C, injured animals did not exhibit increased jumping one day after injury compared to uninjured controls, indicating that the amputation injury did not result in acute hypersensitivity. Seven days following leg amputation injury however, flies now exhibited significantly more jumps at the normally sub-noxious temperature of 38° C compared to uninjured sham controls (Figure 1A-C). This hypersensitivity to 38° C is indicative of thermal allodynia and suggests the development of central sensitization over the seven day time period following the leg amputation injury. Jumping of injured animals at 38°C was prevented by sensory neuron-specific TrpA1 suppression (Figure 1C), demonstrating that the hypersensitivity of injured animals is a thermal nociceptive response.

**Figure 1.**
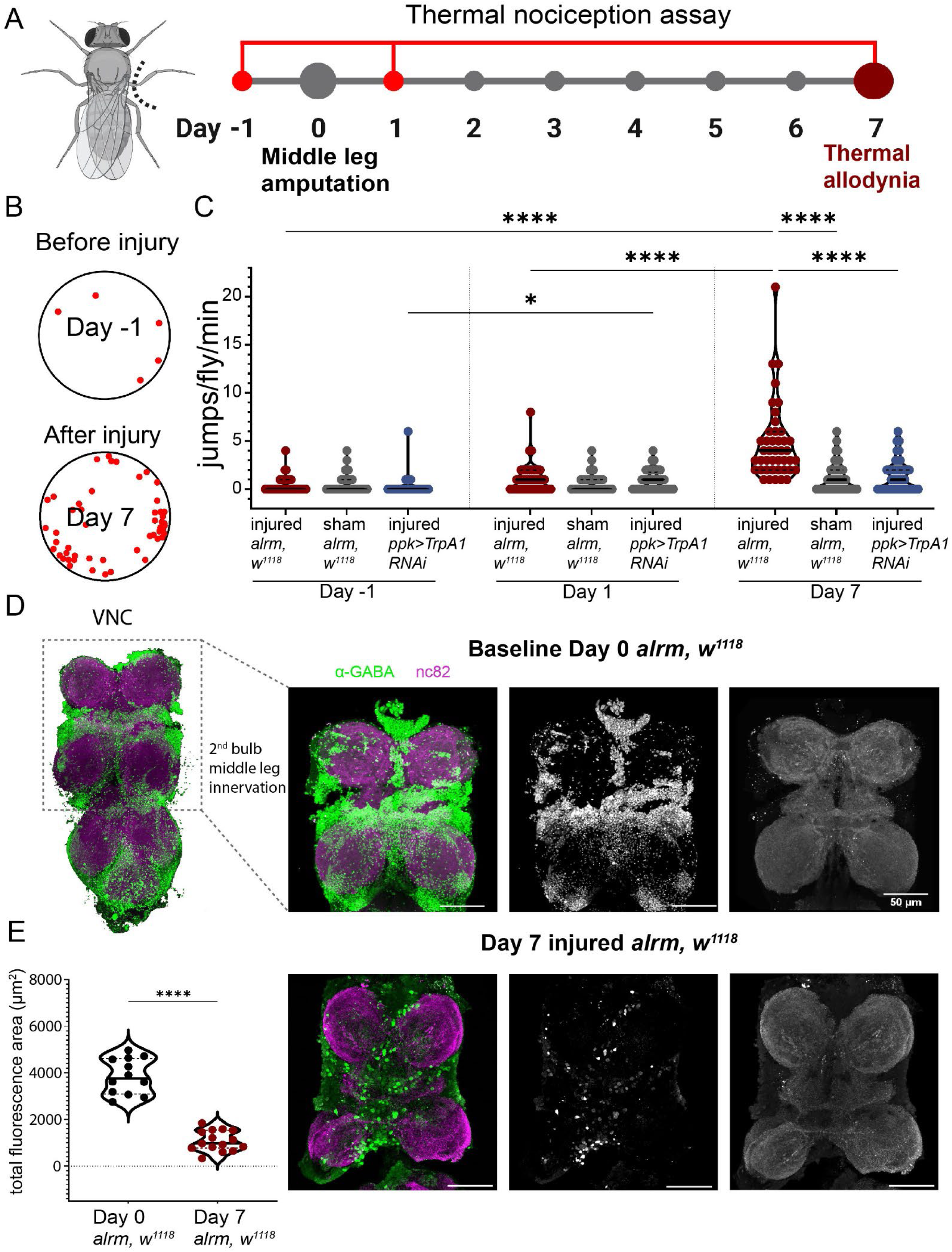
Leg amputation drives centrally-mediated thermal allodynia in adult Drosophila. **(A)** Left: Schematic illustration of nerve injury via middle leg amputation of adult Drosophila. Right: Timeline of behavioral experiments to assess thermal allodynia. As indicated by the red symbols, the same cohorts of animals were tested one day prior to injury (day -1), and one and seven days following injury. **(B)** The same cohort of 11 flies were imaged for one minute at 38° C one day prior to and seven days following injury, and each jump executed by a fly is indicated by a red dot. Shown is a summary diagram of one minute of jumps, represented as red dots, of a group of 11 cohoused flies before injury and seven days after middle leg amputation in an overhead view of a circular chamber on a 38° C hotplate. **(C)** Plot of the number of jumps per fly per minute at 38° C for three cohorts of animals on the indicated days respective to injury. The three cohorts are injured controls (w1118; alrm-GAL4), sham controls (w1118; alrm-GAL4), and injured with *ppk-GAL4* driven *TrpA1* RNAi. Each symbol represents the total number of jumps for a single animal (n≥43). *0.01<P<0.05; ****P<0.0001, Kruskal-Wallis with Dunn’s multiple comparisons, N=411, H(8)=131.7, P<0.0001. **(D)** Representative images of VNCs from uninjured or day seven injured animals labeled with antibodies to α-GABA (green) and bruchpilot (magenta) to label synaptic active zones within the neuropil. **(E)** Plot of the total anti-α-GABA fluorescence area from VNCs of uninjured (n=12) or day seven injured (n=15) animals labeled and imaged as shown in (D). Each symbol represents a single VNC ****, P<0.0001, 2-tailed t-test. Solid lines in violin plots represent medians and dashed lines represent quartiles.

Sensory neurons of the middle legs of flies project onto the second bulbs of the VNC, and it was previously shown that inhibitory GABAergic neuron loss within the second VNC bulbs is both necessary and sufficient for thermal allodynia in response to middle leg amputation (Khuong et al, 2019). Consistent with this, we observed a significant reduction in GABA positive cell bodies in the VNC seven days following leg amputation injury compared to sham controls (Figure 1D). These results collectively establish a robust model of central sensitization and neuropathic pain in adult *Drosophila*, whereby single leg amputation results in thermal allodynia that is accompanied by neural circuitry alterations in the CNS.

### SOCE-dependent astrocyte Ca^2+^ signaling is activated following leg amputation injury

Nerve injury results in enhanced Ca^2+^ wave propagation in astrocytes (Kim et al., 2016; Xu et al., 2021), and complete inhibition of Ca^2+^ responses in cortical astrocytes with the Ca^2+^ chelator BAPTA nearly completely inhibited the development of mechanical allodynia in response to sciatic nerve ligation in mice (Kim *et al*., 2016). However, mechanical allodynia was only partially suppressed in mice lacking expression of the astrocyte-specific type-2 IP_3_R, which mediates Ca^2+^ release from ER Ca^2+^ stores, suggesting the involvement of other astrocyte Ca^2+^ channels in addition to IP_3_R in chronic pain development. To identify additional astrocyte Ca^2+^ signaling mechanisms required for pain sensitization in our *Drosophila* model, we first determined whether leg amputation results in increased Ca^2+^ transients in astrocytes using the Transcriptional Reporter of Intracellular Ca^2+^ (TRIC) expressed with the astrocyte-specific *alrm-GAL4* driver (Doherty et al., 2009). TRIC couples GFP expression to intracellular Ca^2+^ transient activity, and acts as a Ca^2+^ “memory” indicator that provides a quantitative readout of Ca^2+^ signals that occur over a 12-24 hour time period (Gao et al., 2015).

To determine if and when following injury Ca^2+^ signals occur in VNC astrocytes, we imaged GFP fluorescence in the VNC of animals that expressed TRIC transgenes specifically in astrocytes on each of the seven days following leg amputation. Astrocytes also expressed membrane-bound RFP to label astrocytes independently of Ca^2+^ activity, allowing us to calculate a ratio of GFP to RFP fluorescence and factor out Ca^2+^-independent changes to astrocyte fluorescence. Astrocyte GFP fluorescence and the GFP/RFP ratio remained stably low from day zero (prior to injury) until day four after injury, when a highly reproducible spike in astrocyte GFP fluorescence and GFP/RFP ratio was seen throughout the entire VNC (Figure 2A, B). This spike was confined to day four and returned to near-day zero levels from days five through seven. These results suggest that leg amputation injury results in activation of Ca^2+^ signaling events throughout the entire VNC astrocyte network during the three to four day timeframe post-injury.

**Figure 2.**
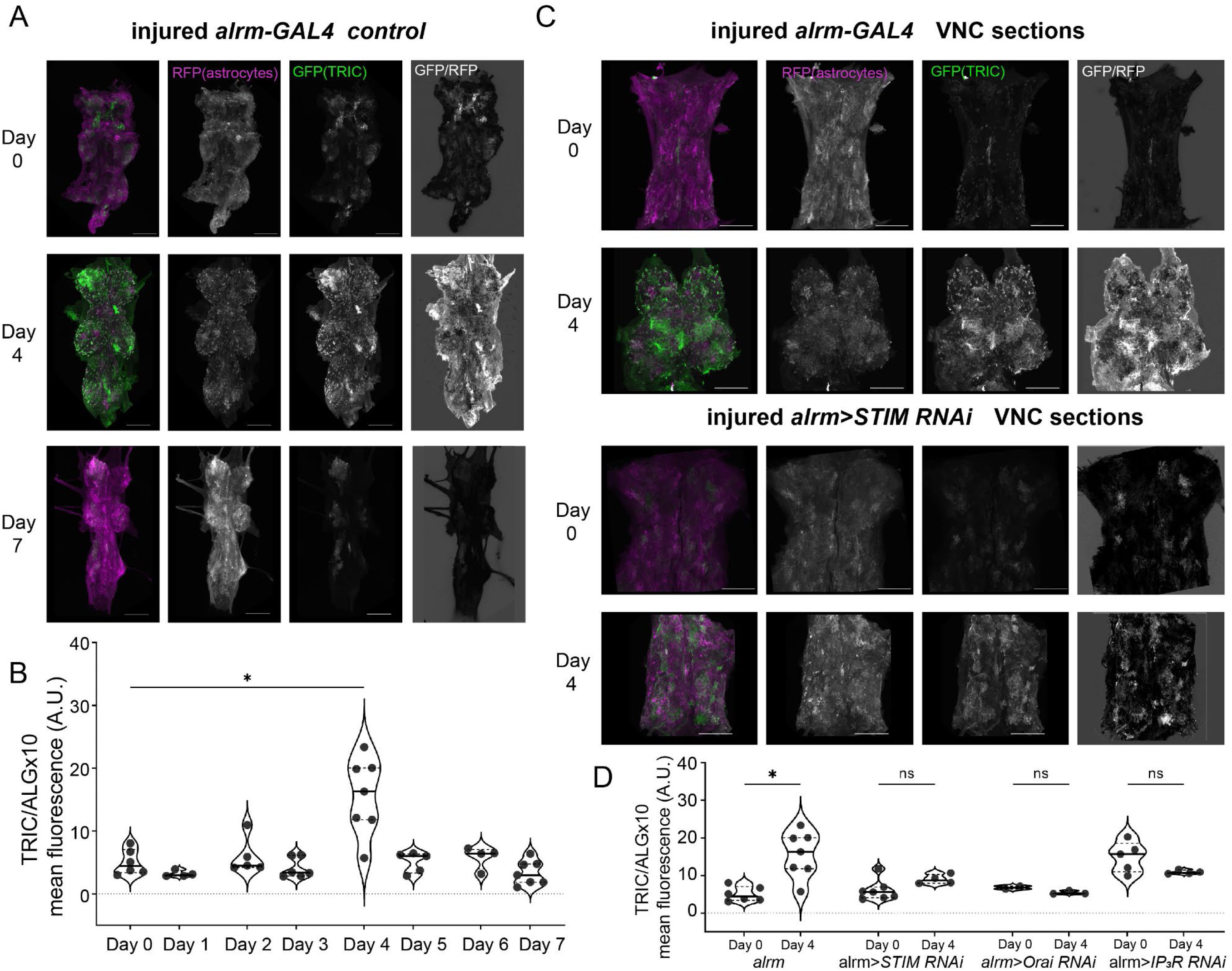
**Leg amputation injury activates SOCE-dependent Ca^2+^ signaling in VNC astrocytes. (A)** Representative images of TRIC-mediated GFP fluorescence (green) and astrocyte-restricted RFP fluorescence (magenta) in VNCs from uninjured (day 0) and day four and day seven post-leg amputation injury animals. Also shown are images of the product of GFP divided by RFP fluorescence, representative of the normalized Ca^2+^ signaling activity reported by TRIC. **(B)** Plot of the mean GFP/RFP fluorescence ratio from VNCs from animals on each day following injury up to day seven. Each symbol represents a single VNC (*n≥4* for each day). *, *P<0.05*, Welch’s ANOVA and multiple comparisons with Dunnett’s T3 correction, *N=44, W(7.000,14.47)=5.100, P=0.0043*. **(C)** Representative images of VNCs cropped to show the anterior two bulbs from seven-day injured control (*w^1118^; alrm-GAL4*) and *alrm-GAL4>Stim RNAi* animals as described in (A). **(D)** Plot of the mean GFP/RFP fluorescence ratio from seven-day control (*w^1118^; alrm-GAL4*) and *alrm-GAL4* driven *Stim*, *Orai*, and *IP_3_R* RNAi VNCs. Each symbol represents a single VNC. *, *P<0.05*, Welch’s ANOVA and multiple comparisons with Dunnett’s T3 correction, *N=33*, *W(5.000, 11.54)=12.83, P=0.0002*; ns, not significant. Solid lines in violin plots represent medians and dashed lines represent quartiles.

We next sought to identify specific Ca^2+^ channels that mediate these astrocyte Ca^2+^ signaling events post-injury. Consistent with prior findings, we found that *IP_3_R* expression was required because astrocyte-specific *IP_3_R* RNAi prevented the day four post-injury spike in TRIC-mediated astrocyte GFP expression (Figure 2D and Supplementary Figure S2). The requirement for IP_3_R-mediated ER Ca^2+^ release suggests the involvement of the SOCE pathway, as SOCE is activated by the depletion of ER Ca^2+^ stores. SOCE requires Stim proteins that function as ER Ca^2+^ sensors and Orai Ca^2+^ influx channels in the plasma membrane. Astrocyte-specific *Stim* and *Orai* suppression using previously validated RNAi constructs (Petersen et al., 2020) also completely prevented the day four spike in TRIC-mediated GFP expression in astrocytes, suggesting that post-injury activation of astrocyte Ca^2+^ signaling requires both IP_3_R-mediated ER Ca^2+^ release as well as Ca^2+^ influx mediated by SOCE.

### Suppression of astrocyte store-operated Ca^2+^ entry attenuates thermal allodynia after leg nerve injury

Suppression of injury-induced astrocyte Ca^2+^ responses by knockdown of Stim and Orai suggests that SOCE mediated by these proteins may also be required for the development of centrally-mediated thermal allodynia. We tested this with repeated thermal nociception assays as described in Figure 1 in animals expressing *Stim* and *Orai* RNAi in astrocytes, as well as *IP_3_R* RNAi as a positive control for an astrocyte Ca^2+^ channel known to be required for allodynia development (Kim *et al*., 2016). Prior to injury, jumps at 24° C and 38° C in astrocyte *Stim*, *Orai*, and *IP_3_R* RNAi groups were relatively few and no different from controls (Supplemental Figure S3A and B), indicating that suppression of these targets alone does not alter jumping behavior. Seven days following leg amputation injury, control RNAi animals exhibited significantly more jumps at 38° C compared to uninjured controls as expected (Figure 3A), indicative of thermal allodynia. In striking contrast, however, jumping at 38° C seven days following injury in animals with astrocyte-specific *Stim*, *Orai*, and *IP_3_R* RNAi was nearly completely suppressed and was no different from uninjured controls (Figure 3A), suggesting inhibition of thermal allodynia in these animals. Suppression of jumping was not due to an overall block of thermal nociception because uninjured *Stim*, *Orai*, and *IP_3_R* RNAi animals exhibited frequent jumps at 42° C similar to controls (Supplemental Figure S3C). Suppression of thermal allodynia was also specific to Stim, Orai, and IP_3_R-mediated Ca^2+^ signaling and not generalizable to other Ca^2+^ channels with known functional activity in astrocytes such as TrpA1 (Shigetomi et al, 2016), as astrocyte suppression of TrpA1 did not prevent thermal allodynia (Supplemental Figure S3D).

**Figure 3.**
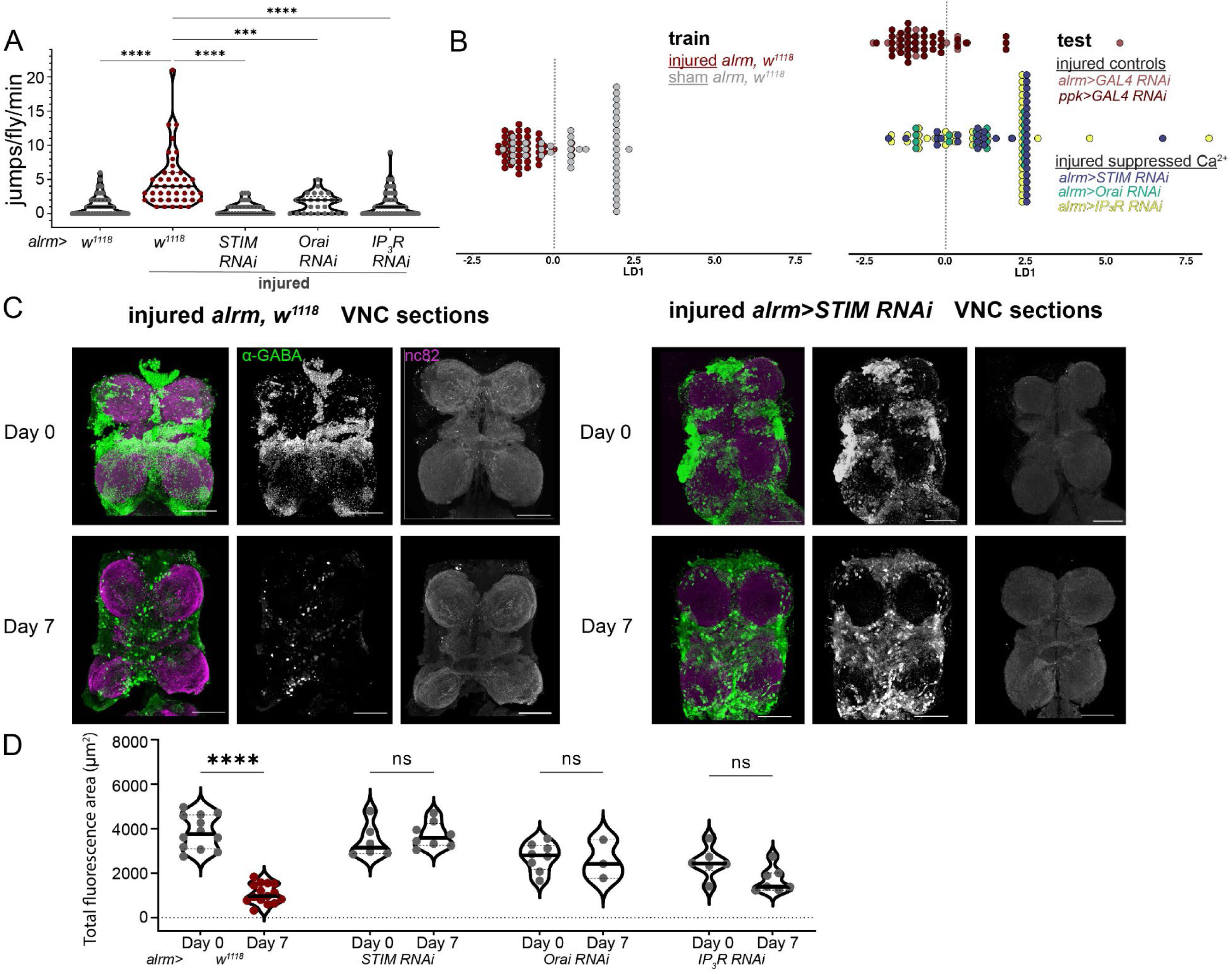
**Suppression of astrocyte SOCE attenuates injury-induced thermal hypersensitivity and GABAergic neuron loss. (A)** Plot of the number of jumps per fly per minute at 38° C for seven days following leg amputation injury (“injured”) for the indicated genotypes. Also shown are data for uninjured controls (left-most dataset). Each symbol represents the total number of jumps for a single animal (N=45 for uninjured *alrm>w^1118^*controls, 43 for injured *alrm>w^1118^* controls, 38 for *Stim RNAi*, 25 for *Orai RNAi*, and 45 for *IP_3_R RNAi*). \*\*\*\**P<0.0001,* \*\*\**P=0.0001*, Kruskal-Wallis with Dunn’s multiple comparisons, *N=196, H(4)=62.94, P<0.0001*. **(B)** Cepstral linear discriminant analysis based on graded behavior over one minute at 38° C for uninjured and seven-day injured animals with indicated genotypes. The left plot shows the result of training the model using data from sham and injured *alrm>w^1118^* animals. The dotted line presents the linear decision boundary, and each symbol represents the linear discriminant or “decision” of the model for each animal based on the deconvolved time series for that animal (classification rate = 0.80). Note that the sham animals cluster to the right of the decision boundary while the injured animals cluster to the left, defining right as uninjured behavior and left as injured behavior. The right plot shows results using test data from two groups of injured controls expressing non-targeting RNAi (*alrm>GAL4 RNAi* and *ppk>GAL4 RNAi*) and injured animals with astrocyte expression of *Stim, Orai,* and *IP_3_R* RNAi. Note that the *Stim*, *Orai*, and *IP_3_R* RNAi animals cluster to the right of the decision boundary consistent with uninjured behavior, despite these animals being injured. **(C)** Representative images of VNCs from uninjured (day 0) or day seven injured control (*alrm; w^1118^*; left) and *alrm>Stim RNAi* (right) animals labeled with antibodies to α-GABA (green) and Bruchpilot (magenta) to label synaptic active zones within the neuropil. **(D)** Plot of the total anti-α-GABA fluorescence area from VNCs of uninjured (day 0) or day seven injured animals with the indicated genotypes. Each symbol represents a single VNC. ****, *P<0.0001*; ns, not significant; mixed effects analysis, multiple comparisons with Šídák’s correction, time *F(1, 57)=24.31*, group *F(2.269, 43.12)=13.94*, and time x group *F(3, 57)=22.03* all *P<0.0001*. Solid lines in violin plots represent medians and dashed lines represent quartiles.

We found in our thermal allodynia assays that fly behavior at 38° C is more complex over the one-minute time series than a simple count of jump number can reflect. For example, some jumps may be frequent and clustered in a short period of time, or they may be spread out over the entire minute, and some flies exhibit responses more severe than jumping such as rolling or seizure-like activity. To account for this complexity, we reanalyzed the videos used for jump counts of seven-day injured flies in Figure 3A by binning the behavior of each fly, per frame of video, into three graded categories to form time series. In this grading scheme, one was assigned to all non-escape or non-avoidance activities (e.g., walking, grooming), two was assigned to jumping, and three was assigned to rolling, seizure-like responses (Supplemental Figure S4A). Using cepstral Fisher’s discriminant analysis (Krafty, 2016), we then trained a discriminant model using time series data from injured and sham *alrm-GAL4, w^1118^* controls. The linear discriminant model performs dimension reduction, extracting characteristics of frequency intervals to plot each fly’s behavior as a single point, and finds a decision boundary that best separates the two given training groups of sham and injured flies. This yielded a highly accurate 0.80 classification rate of injured or sham animals to the injured or sham groups, respectively (Figure 3B). In Figure 3B each fly’s estimated discriminant is plotted along the first linear discriminant (LD1) axis with the linear decision boundary marked as a dotted line. The model was tested using five injured fly groups. It classified a majority of the two new control injured fly groups as behaving like injured flies, whereas 81.58% of *Stim* RNAi, 70.83% of *Orai* RNAi, and 68.89% of *IP_3_R* RNAi injured flies were predicted to be sham (Figure 3B and Supplemental Figure S4C). This further validates that even when taking into account behavior patterns more complex than jump number, astrocyte-specific Stim, Orai, and IP_3_R suppression prevents behaviors associated with thermal allodynia following injury.

We next determined whether GABAergic neuron loss following leg amputation injury requires Stim, Orai, and IP_3_R function in astrocytes. Consistent with this, we found that GABA immunoreactivity in VNCs from animals with astrocyte-specific *Stim*, *Orai*, or *IP_3_R* RNAi was unchanged and remained high at day seven following injury compared to day zero (uninjured), in clear contrast to injured controls that showed a significant reduction in GABA on day seven post-injury (Figure 3C and D and Supplemental Figure S5). These results suggest that Ca^2+^ signaling mediated by Stim, Orai, and IP_3_R is required in astrocytes for neural circuitry changes in the CNS that result in thermal allodynia.

### Astrocyte SOCE is sufficient for development of thermal allodynia

Our results thus far suggest that astrocyte SOCE function is required for development of thermal allodynia following leg amputation injury in flies. We next asked whether astrocyte SOCE is sufficient for thermal allodynia. We tested this by expressing in astrocytes a constitutively active Orai mutant that has a glycine to methionine substitution (Orai^G170M^) in the hinge region of the channel that forces the channel into an open conformation (Zheng et al., 2013). TRIC-mediated GFP expression was significantly higher in astrocytes from uninjured animals with astrocyte-specific Orai^G170M^ expression compared to non-expressing controls (Figure 4B, C), suggesting that Orai^G170M^ expression increases astrocyte Ca^2+^ signaling in the absence of injury as expected. Strikingly, animals with astrocyte-specific Orai^G170M^ expression exhibited thermal allodynia even when uninjured, as their number of jumps at 38° C was similar to seven-day injured, non-expressing controls and significantly greater than uninjured controls (Figure 4A). This significant increase in jumping was not seen in uninjured animals expressing a wildtype *Orai* transgene (*Orai^+^*), suggesting that astrocyte Ca^2+^ influx is specifically responsible for the development of hypersensitivity. Our results collectively demonstrate that astrocyte SOCE is both necessary and sufficient for the development of thermal allodynia in adult *Drosophila*, and that astrocyte Ca^2+^ signaling mediated by SOCE is an essential component of the CNS adaptations that underlie the transition from acute nerve injury to neuropathic hypersensitivity.

**Figure 4.**
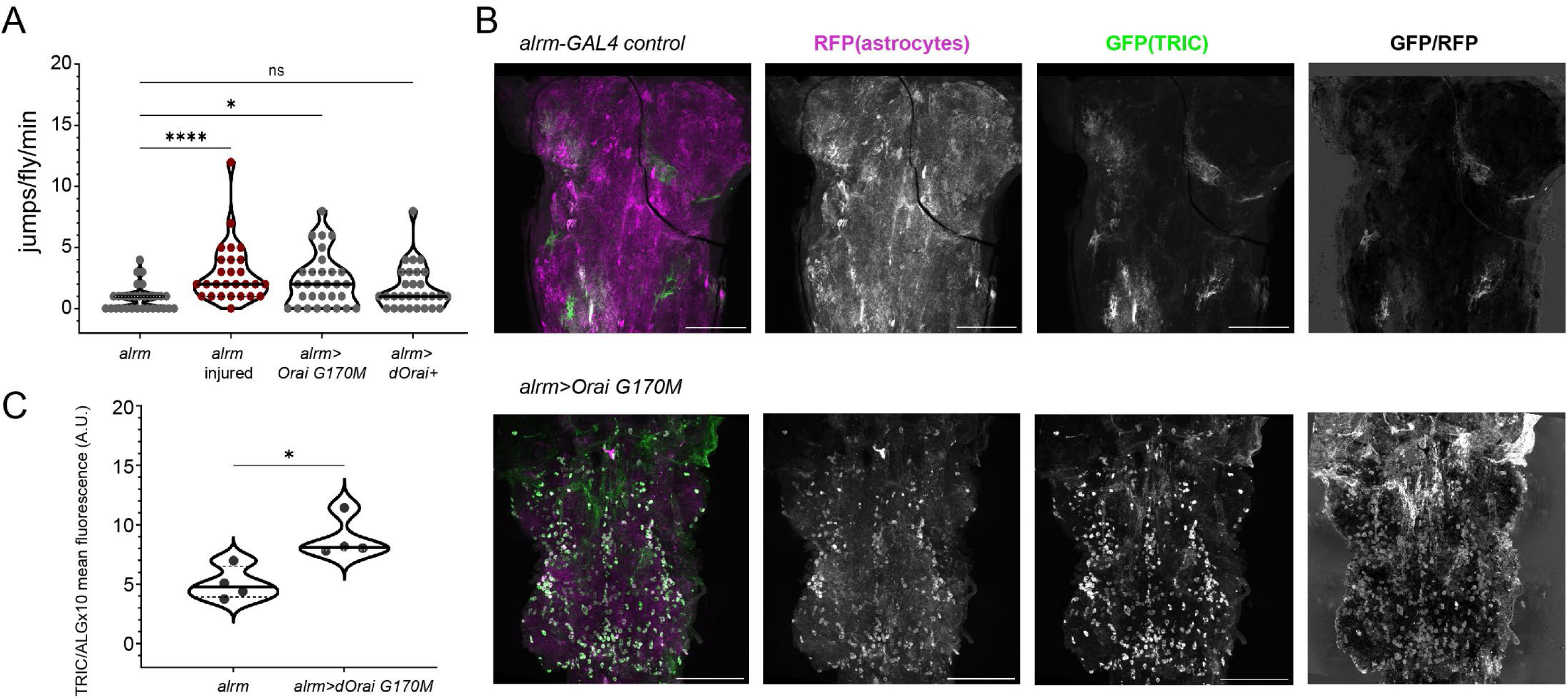
**Constitutive SOCE is sufficient to cause thermal hypersensitivity in the absence of injury. (A)** Plot of the number of jumps per fly per minute at 38° C for uninjured animals with the indicated genotypes, as well as day seven injured controls. Each symbol represents the total number of jumps for a single animal (n=30 for *alrm; w^1118^*, 30 for *dOrai^G170M^*, 29 for *dOrai^+^*, and 27 for injured *alrm; w^1118^*), * *0.01<P<0.05*; ***, *P<0.0001;* ns, not significant; Kruskal-Wallis with Dunn’s multiple comparisons, *N=116, H(3)=19.57, P=0.0002*. **(B)** Representative images of TRIC-mediated GFP fluorescence (green) and astrocyte-restricted RFP fluorescence (magenta) in VNCs from uninjured control (*alrm; w^1118^*) and uninjured *alrm>dOrai^G170M^*animals. Also shown are images of the product of GFP divided by RFP fluorescence, representative of the normalized Ca^2+^ signaling activity reported by TRIC. **(C)** Plot of the mean GFP/RFP fluorescence ratio from uninjured control (*alrm; w^1118^*) and *alrm-dOrai^G170M^* VNCs. Each symbol represents a single VNC. *\*, P<0.05*, Two-tailed Mann-Whitney U Test, *U=0, P=0.0286*. Solid lines in violin plots represent medians and dashed lines represent quartiles.

## Discussion

It has long been recognized that Ca^2+^ signaling is a key mechanism by which astrocytes respond to and in turn modulate neural activity. Astrocyte Ca^2+^ signaling is also augmented following many forms of neurotrauma and nerve injury. Yet despite the central role of Ca^2+^ signaling in overall astrocyte physiology and neuropathology, the specific Ca^2+^ signaling mechanisms and Ca^2+^ channels that regulate astrocyte functions are not well understood. Recent findings demonstrate that Ca^2+^ influx through SOCE channels formed by Orai1 in mouse astrocytes drives gliotransmitter release and tonic inhibition of pyramidal CA1 neurons, clearly defining a role for astrocyte SOCE in physiological neural circuitry and function (Toth et al., 2019). Our results in *Drosophila* now demonstrate for the first time that astrocyte SOCE is also activated in response to neurotrauma and is essential for the neural circuitry changes that underlie central sensitization and chronic neuropathic pain. SOCE may therefore play fundamental and highly conserved roles in multiple aspects of astrocyte physiology.

It is notable that our TRIC Ca^2+^ assays revealed a distinct and relatively narrow period of enhanced astrocyte Ca^2+^ signaling three to four days following injury. This is in fact consistent with data from mice showing that partial sciatic nerve ligation results in a significant increase in cortical astrocyte Ca^2+^ transient frequency between three and six days post-injury, with a return to basal frequency after day six (Kim *et al*., 2016). The timing of allodynia development is also similar between our *Drosophila* model and mouse models, with maximal allodynia at six to nine days post-injury and persistence for several weeks (Kim *et al*., 2016; Kim and Nabekura, 2011). Thus, enhanced astrocyte Ca^2+^ signaling may be a relatively early and temporally confined response during the progression from acute injury to neuropathic pain. We also found that astrocyte Ca^2+^ responses indicated by TRIC were not confined to the second VNC bulbs where the sensory nerves from the injured leg innervate, but instead were distributed throughout the VNC. It is possible that gap junctions between astrocytes are responsible for Ca^2+^ signals that occur in astrocytes that are distant from the site of innervation of the damaged nerves. It is well documented that astrocytes form highly interconnected networks throughout the CNS via gap junctions that propagate signals including Ca^2+^ (Panattoni et al., 2021; Pasti et al., 1997), and astrocyte gap junctions are required for neuropathic pain following acute injury in mice (Chen et al., 2012). Consistent with this, we found that astrocyte-specific suppression of the *Drosophila* gap junction component innexin-2 partially suppressed thermal allodynia following leg amputation injury (Supplemental Figure S6).

Release of ER Ca^2+^ by IP_3_Rs in astrocytes is clearly important for neuropathic pain based on our current results and prior findings in mice (Kim *et al*., 2016). IP_3_R-mediated Ca^2+^ release results in “somatic” Ca^2+^ signals in astrocytes that occur throughout the cell body near the centrally-located nucleus (Bazargani and Attwell, 2016). It was long thought that these IP_3_R-mediated Ca^2+^ responses, which occur downstream of metabotropic receptor activation, constitute the primary mechanism by which astrocytes respond to synaptic activity. However, it is now becoming increasingly clear that “microdomain” Ca^2+^ signals that are confined to thin astrocytic processes are also essential components of Ca^2+^-dependent astrocyte information processing (Arizono et al., 2020; Bindocci et al., 2017; Otsu et al., 2015; Rungta et al., 2016; Stobart et al., 2018). Our major finding that astrocyte SOCE is essential for neuropathic pain is highly significant in this regard, because SOCE produces localized, microdomain Ca^2+^ signals that function independently of IP_3_R-mediated Ca^2+^ release (Di Capite et al., 2009; Parekh, 2011). It is likely that SOCE in astrocytes is activated subsequent to IP_3_R-mediated ER Ca^2+^ release, and the resultant depletion of ER Ca^2+^ stores serves as the signal that activates SOCE. The combination of IP_3_R-mediated somatic Ca^2+^ signals and SOCE-mediated microdomain Ca^2+^ signals may therefore provide a robust, multifaceted Ca^2+^ signaling paradigm downstream of metabotropic receptor activation that results in highly efficient and specific activation of astrocyte functions during the transition from acute injury to neuropathic pain. It has also been shown that Waterwitch (Wtrw) and TrpML Ca^2+^ channels mediate somatic and microdomain astrocyte Ca^2+^ signals in *Drosophila*, respectively (Ma and Freeman, 2020; Ma et al., 2016), and we found that astrocyte-specific suppression of these channels also partially attenuated thermal allodynia in flies (Supplemental Figure S7). It will be interesting to determine in future experiments whether these channels function in coordination with or independently of IP_3_R and SOCE channels in astrocytes during neuropathic pain signaling.

Our findings demonstrate that astrocyte SOCE is required not only for the development of thermal allodynia, but also for the neural circuitry changes that occur subsequent to injury as revealed by GABA immunoreactivity in the VNC. The loss of GABAergic neurons following injury is likely to remove tonic inhibitory signals to nociceptive circuits, resulting in hyperexcitability and hypersensitivity associated with allodynia (Basbaum et al., 2009). Loss of GABAergic neurons is due to apoptosis (Khuong *et al*., 2019), though it is not clear how apoptosis of these neurons is activated. We cannot determine from our experiments whether SOCE-regulated astrocyte function is directly responsible for GABAergic neuron loss, or whether astrocyte function is a component of a cascade of events that results in neuronal apoptosis. A major goal moving forward will be to determine the specific targets of SOCE in astrocytes, and how these targets regulate astrocyte functions that are essential for central sensitization and neuropathic pain. SOCE in cortical mouse astrocytes drives the exocytic release of the gliotransmitters ATP and D-serine (Toth *et al*., 2019), and a similar function of SOCE activated by nerve injury may alter synaptic function within nociceptive circuits. Astrocytes also secrete inflammatory cytokines in response to Ca^2+^ signals (Hennessy et al., 2015; Liddelow et al., 2017), and cytokines may also modulate neuronal function or could induce GABAergic neuron apoptosis. Interestingly, it was recently shown in mice that nerve injury-induced SOCE in microglia is required for neuropathic pain due to SOCE-dependent microglial cytokine production (Tsujikawa et al., 2023). While flies do not have the equivalent of microglia within their CNS, SOCE-mediated cytokine release may be a major glial cell function in the pathogenesis of neuropathic pain.

In conclusion, our results demonstrate that astrocyte SOCE function is an essential component of the neural adaptations that lead to central sensitization and neuropathic pain following acute nerve injury. These findings may drive the development of new therapeutics for neuropathic pain that do not rely on highly addictive and dangerous drugs such as opioids. In addition, our findings further demonstrate that SOCE is an essential and highly conserved component of astrocyte function in neural physiology and pathogenesis. This will contribute significantly to our understanding of the complexity and diversity of Ca^2+^ signaling processes in astrocytes.

## Methods

### Materials

A table listing all materials and reagents with source information used in this study is included in Supplementary Materials (Supplemental Table 1).

### Fly genetics & husbandry

*Drosophila* were housed in vials on fly food (yeast: 10 g; corn syrup: 65 g; corn meal: 42.5 g; soy flour: 5.625 g; propionic acid: 4.25 mL/L, agar 5.125 g; 95% tegocept: 15 mL, H_2_O: 0.8 L). Experimental groups were separated into vials by sex in groups of 10-12. Flies were 5-8 days old at the time of leg amputation injury. To ensure consistent baseline locomotion, fly lines were raised and kept at 25 °C with a 12-hour light/dark cycle (Giraldo et al., 2019; Wheeler et al., 1993). Leg amputations, behavioral experiments, and dissections were conducted at the same morning timepoints. For the longer duration thermal nociception assay, all males were tested first to account for circadian activity cycles (Helfrich-Förster, 2000). In order to incorporate all the required genetic components (e.g. drivers, reporters, RNAis, mutations), recombination was conducted over four generations using balanced stocks and tracking elements on both second and third chromosomes (Prokop, 2013).

### Thermal allodynia assay

Thermal nociception assays (Khuong *et al*., 2019) were conducted by placing a group of 10-12 flies in a round chamber 3 mm high with a 35 mm diameter (Vellum translucent paper glued to a 3D printed ring) on a heat block (Benchmark Scientific) set to 38°, 24°, or 42° C and after 60 sec of acclimation, fly behavior was recorded for 90 sec at 30 frames/sec on a Google Pixel smartphone from above. The heat block was calibrated and checked between groups using an infrared thermometer gun (Fisher Scientific). Videos were cropped in size and duration and converted to AVI files using MATLAB, and jumping behavior was visually scored as described below.

For the thermal allodynia experiment, each experimental genotype was divided into 4 replicate groups (2 male, 2 female) of 10-12 flies to be injured and 4 replicate groups to be sham. The 38° C experiment was conducted on Day -1, 1, and 7 relative to injury. On Day 0, flies designated as injured had the right middle leg amputated mid-femur under CO_2_ anesthesia. Sham flies were placed under CO_2_ anesthesia for the same duration. The experimenter was blinded for behavior collection and analysis.

### Behavioral data analysis

Blinded videos were manually analyzed using FIJI and jump number and the start and stop frames of jumps and rolls were recorded. Due to right skewedness from large number of flies that had zero jumps, all jump number data were analyzed using non-parametric Kruskal-Wallis tests with Dunn’s correction for multiple comparisons. Male and females were initially analyzed separately, but no sex differences were found. Male and female data was therefore pooled for the reported analyses. For the linear discriminant analysis (LDA), each fly’s behavior was coded into a time series based on jump and roll start and stop frames using R. To analyze the time interval relationships between jumps and rolls in every fly, the time series were transformed into frequency domain spectra via fast Fourier transform and then into time domain cepstra via inverse Fourier transform. Using the CepLDA package in R (Krafty, 2016), cepstral Fisher’s LDA was run by first training the model to find the best weights vector to separate sham and injured control flies reacting to 38° C on Day 7, then running the LDA model on new injured test groups.

### Dissection and immunohistochemistry

We used Janelia Flylight dissection protocols for dissection and immunohistochemistry of adult *Drosophila* CNS (Jenett et al., 2012). In brief, prior to dissection flies were placed in a 4° C refrigerator for 10 min in vials. Cold anesthetized flies were individually transferred to a 9-well glass plate (Pyrex) on ice, rinsed in 70% ethanol, twice rinsed in S2 media (Schneider’s Insect Medium, Fisher Scientific), and dissected in cold S2 media on a 9-well glass plate coated with silicone (Sylgard 184, Sigma) using forceps (Dumont #5, Fine Science Tools). We first pulled away the thorax and abdomen from the VNC, then dissected the head and pulled away the brain. Dissected VNCs were transferred into 2% PFA in S2 media and nutated at room temperature (RT) for 55-65 min, then washed four times for 10 min each in phosphate buffered saline (PBS) with 0.5% Triton X-100 (PBT) with nutating at RT. PBT was removed and VNCs were incubated in blocking buffer (500 mg bovine serum albumin, 10 mL PBT) for 1 hr with nutating at RT, then transferred to primary antibody solution and nutated first at RT for 4 hr then at 4° C for 48 hours.

All antibodies were diluted in blocking buffer. Primary antibodies used were mouse anti-Bruchpilot nc82-s (DSHB) 1:20, rabbit anti-GABA (Sigma) 1:50, rabbit anti-dsRed (Takara Bio) 1:500, and chicken anti-GFP (Abcam) 1:500. VNCs were then washed four times for 15 min each in PBT with nutating at RT, then transferred into secondary antibody solution and nutated first at RT for 4 hr then at 4° C for 72 hours. Secondary antibodies used were Alexa fluor 568 goat anti-mouse 1:500, Alexa fluor 488 goat anti-rabbit 1:500, Alexa fluor 568 goat anti-rabbit 1:800, and Alexa fluor 488 goat anti-chicken 1:800.

Prior to sample mounting, a cover glass (Corning No. 1, Fisher Scientific) was dipped overnight in PLL solution (Poly-L-lysine hydrobromide powder, 32 mL of diH2O, 64 µL Kodak Photo-Flo) and stored in darkness at 4° C. After secondary antibody incubation, VNCs were rinsed and washed four times as describe, then transferred into 4% PFA in PBS for a secondary fixation with nutating for 4 hr at RT. VNCs were again washed four times in PBT for 15 min and then once more in PBS for 10 min. VNCs were mounted onto PLL-dipped cover glass in PBS, up to 20 samples per cover glass. The cover glass with mounted VNCs was dipped in diH2O, then dehydrated in successive 10 min baths of ethanol: 30%, 50%, 75%, 95%, 100%, 100%, 100%. Finally, drops of DPX (Electron Microscopy Sciences) were added to the cover glass until all the samples were covered, then the cover glass was flipped onto a glass slide with 2 cover glass spacers <20 mm apart, and the slide cured at RT for 3 days before imaging.

### GABA imaging and analysis

Samples were imaged with a Nikon A1R confocal microscope using a 40X, 1.3 NA oil objective. Z-stacks at 0.4 µm intervals were collected. Using MATLAB, image files were blinded using a seeded random integer generator and script to rename files and save the blinding key. Image processing and analysis were performed using Fiji. Images were imported as ND2 stacks, channels were split, and the stacks of GABA immunofluorescence from 488 nm excitation were processed with background subtraction, then stacks were Z-projected showing maximum intensity over the entire depth of the VNC. Because GABA immunofluorescence appeared as punctate, particle-like structures, we determined the total area of GABA particles to quantify GABA immunofluorescence. To accomplish this, image threshold was adjusted to include the maximum fluorescent particles and minimum noise, the image was converted to a binary mask, small noise particles were removed (binary>open command) and holes were filled if they appeared inside particles. Finally, particles were analyzed with the following settings: 0-infinity µm^2^, 0-1.0 circularity, summarize. The summary output total fluorescence area (µm^2^) was reported. To analyze the quantification of GABA immunoreactivity in controls before and after injury, we used a 2-tailed t-test because the data met conditions of normality and equal variances between groups. To analyze changes in GABA after injury in all groups, we used a non-parametric mixed-effects analysis with Šídák’s correction for multiple comparisons because these data did not meet normality or homoscedasticity requirements.

### TRIC imaging and analysis

Samples were imaged with a Nikon A1R confocal microscope using a 60X, 1.4 NA oil objective. Z-stacks at 0.5 µm intervals were collected, with identical acquisition laser settings and pinhole size. Images were saved as maximum intensity projections of Z-stacks, then blinded as described above. In Fiji, maximum projection images were imported and processed using the calculator plus. The 488 nm excitation channel showing TRIC fluorescence was divided by the 568 nm excitation channel showing total astrocyte fluorescence, then multiplied by 10 for easier visualization. A circular region of interest (ROI) covering 58,436 pixels^2^ was used to measure mean intensity at 5 locations per divided image, covering each of the 4 bulbs of the VNC and one central ROI. For each image, the average of the 5 output intensity means is reported. Because the data are normally distributed but show unequal variances between groups, we used Welch’s ANOVA with Dunnett’s T3 correction for multiple comparisons for TRIC quantification analysis.

## Acknowledgements

This work was supported by NIH grant R21 NS121821 to J.T.S. Reagents obtained from the Bloomington *Drosophila* Stock Center (NIH P40OD018537) were used in this study.

## Notes

### Competing Interest Statement

The authors have declared no competing interest.

