## Supplementary Figures and Table for "Astrocyte store-operated calcium entry is required for centrally mediated neuropathic pain"

**Supplementary Material**

#### Supplemental Figure S1

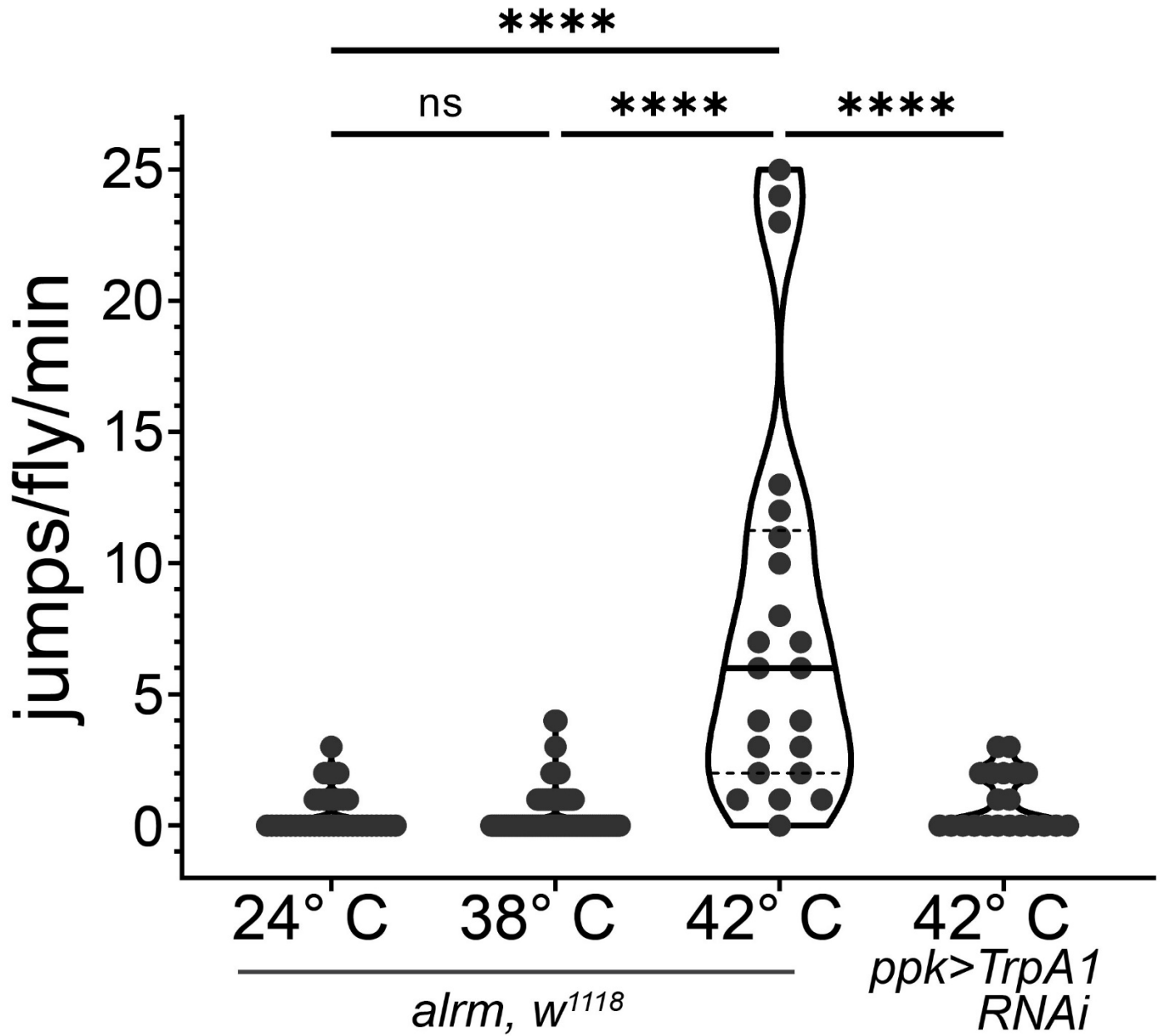

**Supplemental Figure S1. Jumping response is antinociceptive and initiates at a threshold greater than 38° C in healthy flies.** Plot of the number of jumps per fly per minute at 38° C for indicated genotypes and at indicated temperatures. All animals were uninjured. Each symbol represents the total number of jumps for a single animal ($n=42$ for 24°, $n=93$ for 38°, $n=22$ for 42° *alrm-GAL4*; *w<sup>1118</sup>* controls and $n=21$ for *ppk>TrpA1 RNAi*). \*\*\*\*, $P<0.0001$; ns, not significant; Kruskal-Wallis with Dunn's multiple comparisons.

### Supplemental Figure S2

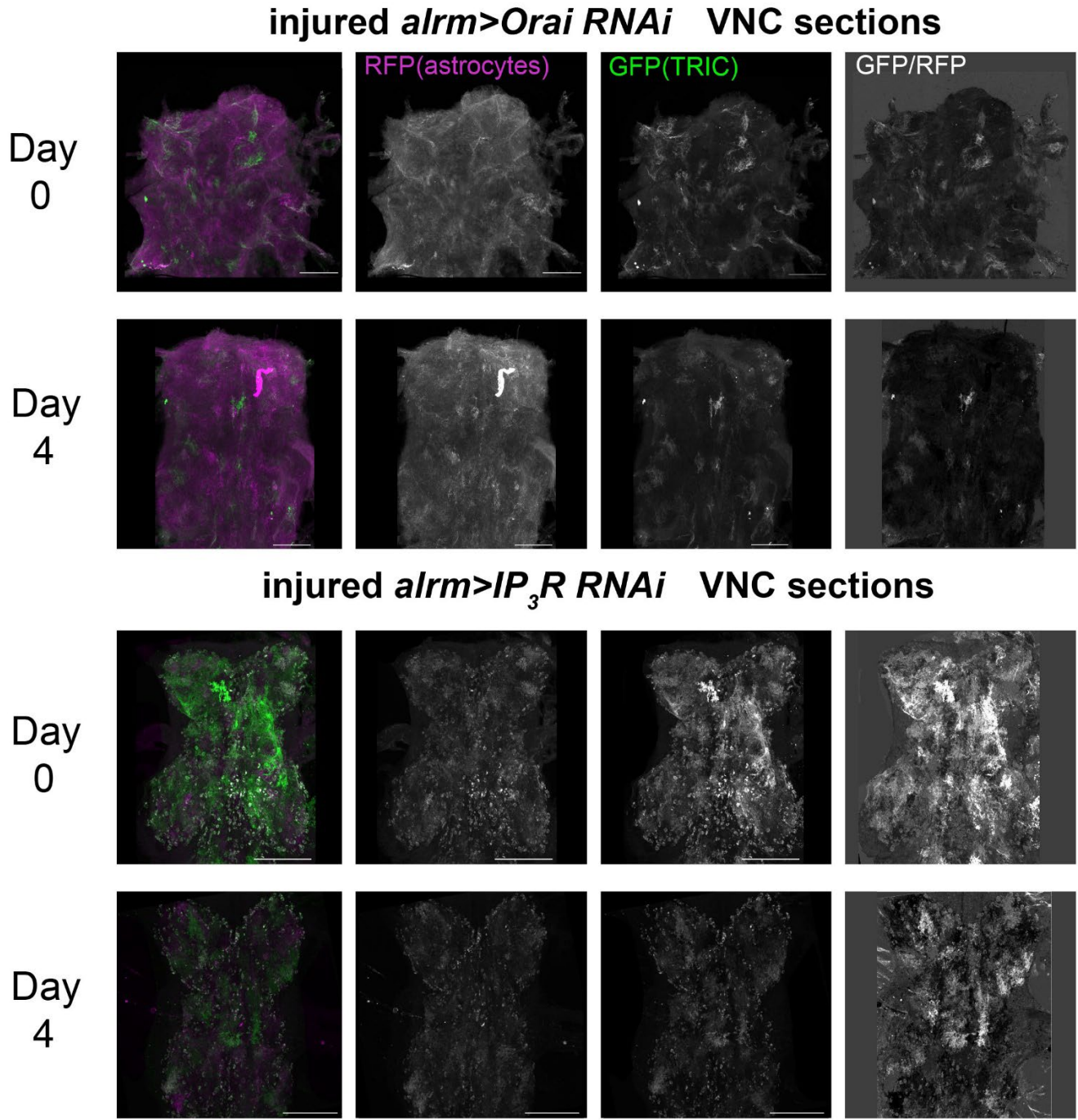

**<u>Supplemental Figure S2.</u>** Astrocyte $\text{Ca}^{2+}$ signaling does not increase four days after injury if $\text{IP}_3\text{R}$ or *Orai* is **suppressed**. Representative images of TRIC-mediated GFP fluorescence (green) and astrocyte-restricted RFP fluorescence (magenta) in VNCs from day 0 (uninjured) and day 4 post-injury *alrm-GAL4>Orai RNAi* and *alrm-GAL4>IP<sub>3</sub>R RNAi* animals. Also shown are images of the product of GFP divided by RFP fluorescence, representative of the normalized $\text{Ca}^{2+}$ signaling activity reported by TRIC.

### Supplemental Figure S3

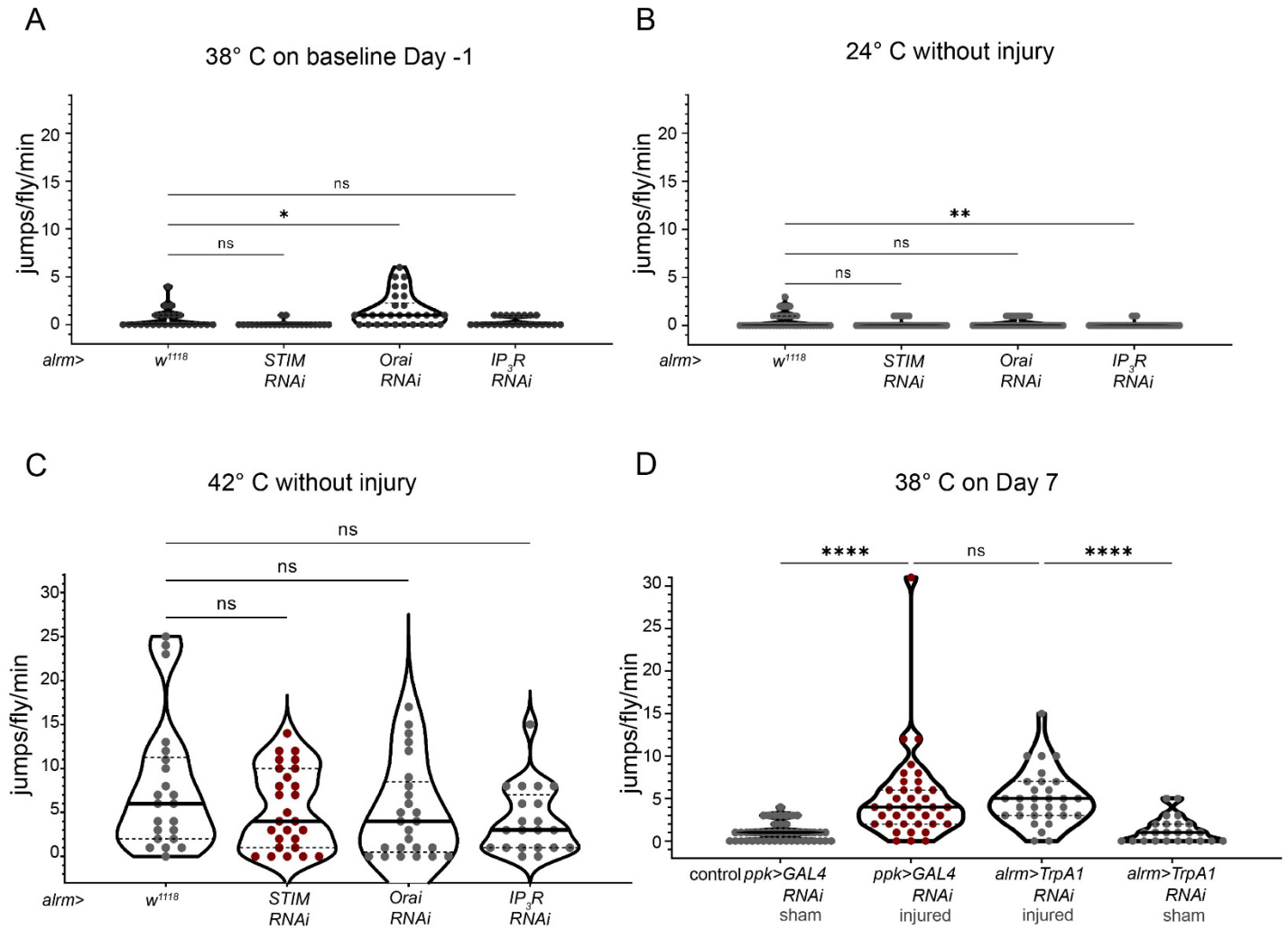

**Supplemental Figure S3. Suppression of astrocyte store-operated calcium entry components does not alter normal jumping activity or nocifensive response to high temperature.** (A) Plot of the number of jumps per fly per minute at 24° C for uninjured flies with indicated genotypes. (B) Plot of the number of jumps per fly per minute at 38° C for uninjured flies with indicated genotypes. (C) Plot of the number of jumps per fly per minute at 42° C for uninjured flies with indicated genotypes. (D) Plot of the number of jumps per fly per minute at 38° C for sham or seven-day injured flies with indicated genotypes. \*0.01 < P < 0.05; \*\*0.001 < P < 0.01; \*\*\*\*P < 0.0001; ns, not significant; Kruskal-Wallis with Dunn's multiple comparisons, N=101, H(3)=22.08, P<0.0001, N=174, H(3)=9.871, P=0.0197, N=94, H(3)=3.267, P=0.3523, N=40, H(3)=58.85, P<0.0001 respectively.

**Supplemental Figure S4**

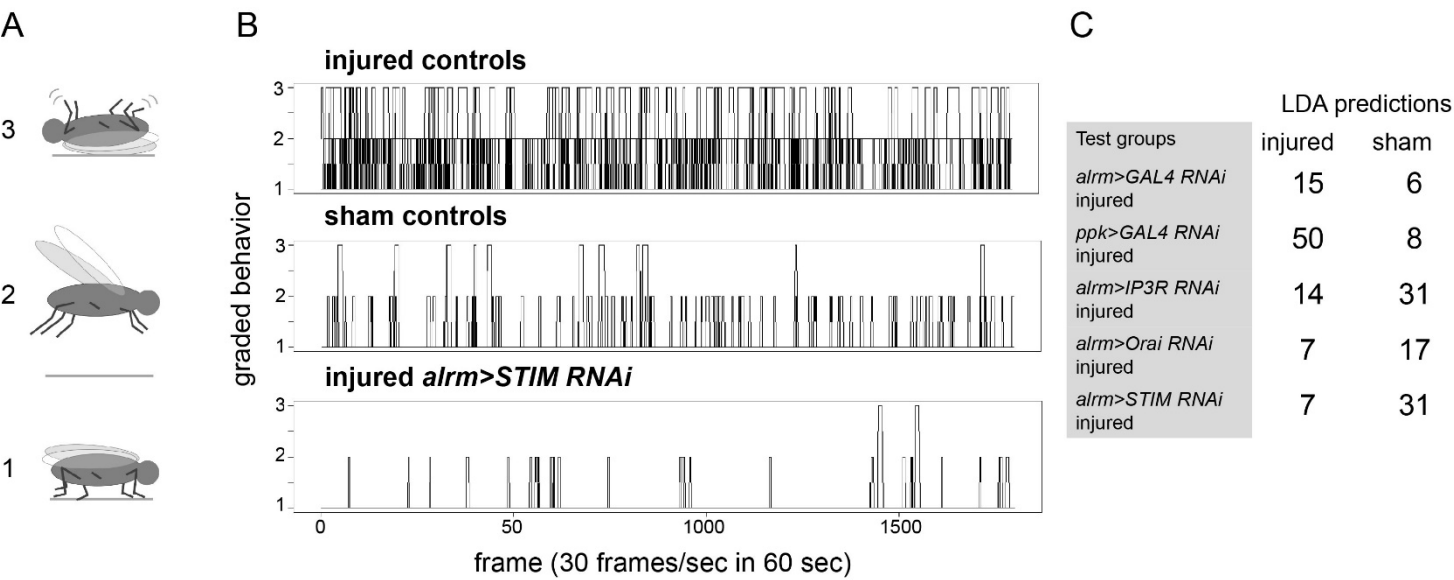

**Supplemental Figure S4. Grading of fly behavior 7 days after injury for cepstral linear discriminant analysis.** (A) For every frame of video (1800 frames in 1 minute at 30 frames per second), each fly’s behavior was binned according to an ordinal scale depending on intensity of reaction to the thermal stimulus. 1=all behaviors not considered anti-nociceptive, 2= jumping, 3=rolling, not able to recover on feet. (B) 1-minute time series generated after binning behavior presented cumulatively per injured controls, sham controls, and STIM RNAi group. Individual fly time series are used as model input with identified conditions (injured or sham) for the training and deidentified for the testing (C) Linear discriminant model predictions from the test groups. All test groups are injured animals; 15/21 *alrm>GAL4 RNAi* and 50/58 *ppk>GAL4 RNAi* controls were categorized correctly as injured, 31/45 *alrm>IP3R RNAi*, 17/24 *alrm>Orai RNAi*, and 31/38 *alrm>STIM RNAi* were predicted to be sham despite injury.

### **Supplemental Figure S5**

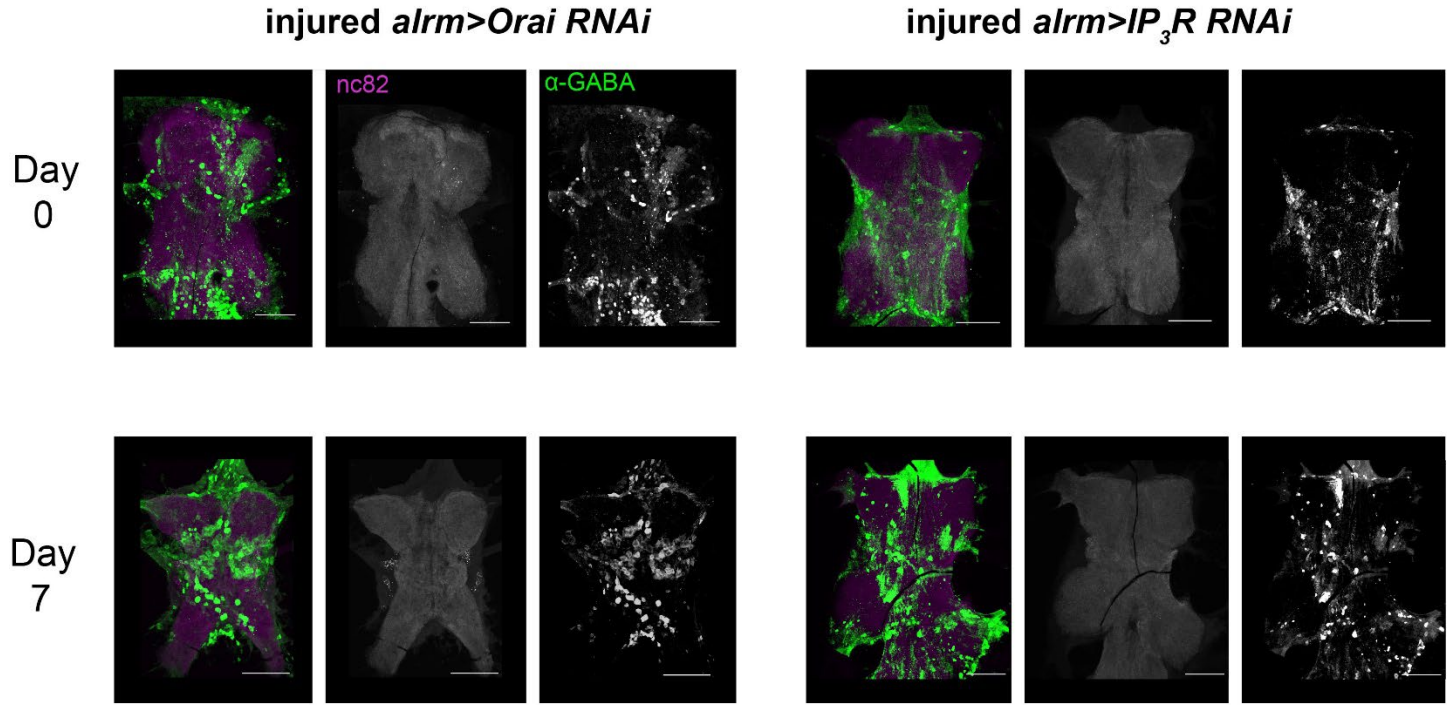

**<u>Supplemental Figure S5.</u>** GABAergic neuron loss following injury is prevented by astrocyte Orai and IP<sub>3</sub>R suppression. Representative images of VNCs from day seven injured *alrm>Orai RNAi* (left) and *alrm>IP<sub>3</sub>R RNAi* (right) animals labeled with antibodies to α-GABA (green) and Bruchpilot (magenta) to label synaptic active zones within the neuropil.

### Supplemental Figure S6

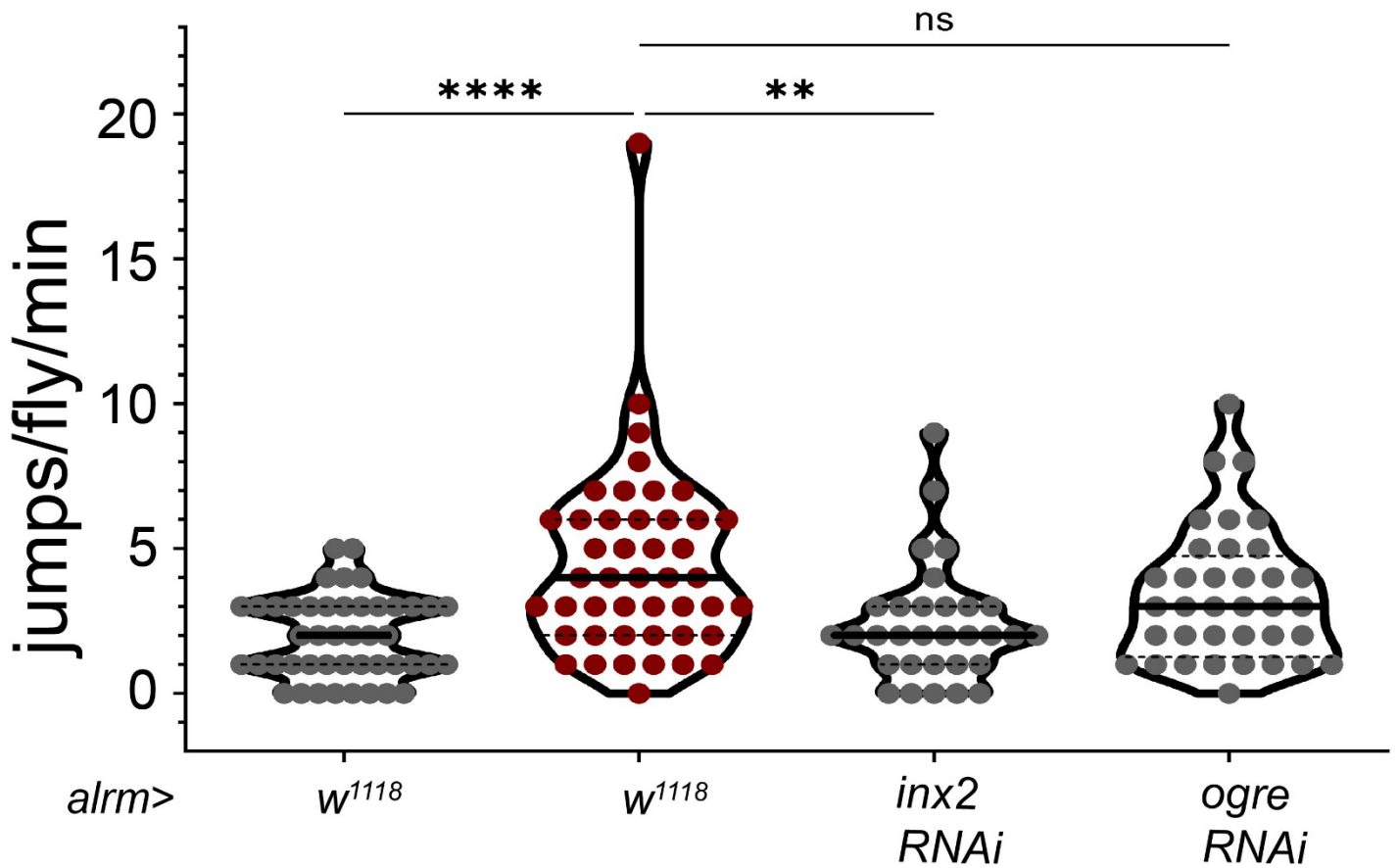

**Supplemental Figure S6. Astrocyte gap junction subunit suppression prevents thermal hypersensitivity following injury.** Plot of the number of jumps per fly per minute at 38° C for seven days following leg amputation injury (“injured”) for the indicated genotypes. Each symbol represents the total number of jumps for a single animal ($n=45$ for *alm-GAL4*; *w<sup>1118</sup>*, $n=31$ for *alrm>inx2 RNAi*, $n=36$ for *alm>ogre RNAi*, and $n=45$ for *alm-GAL4*; *w<sup>1118</sup>* sham controls) \*\*\*\*, $P<0.0001$; Kruskal-Wallis with Dunn’s multiple comparisons, $N=157$, $H(3)=24.63$, $P<0.0001$.

### Supplemental Figure S7

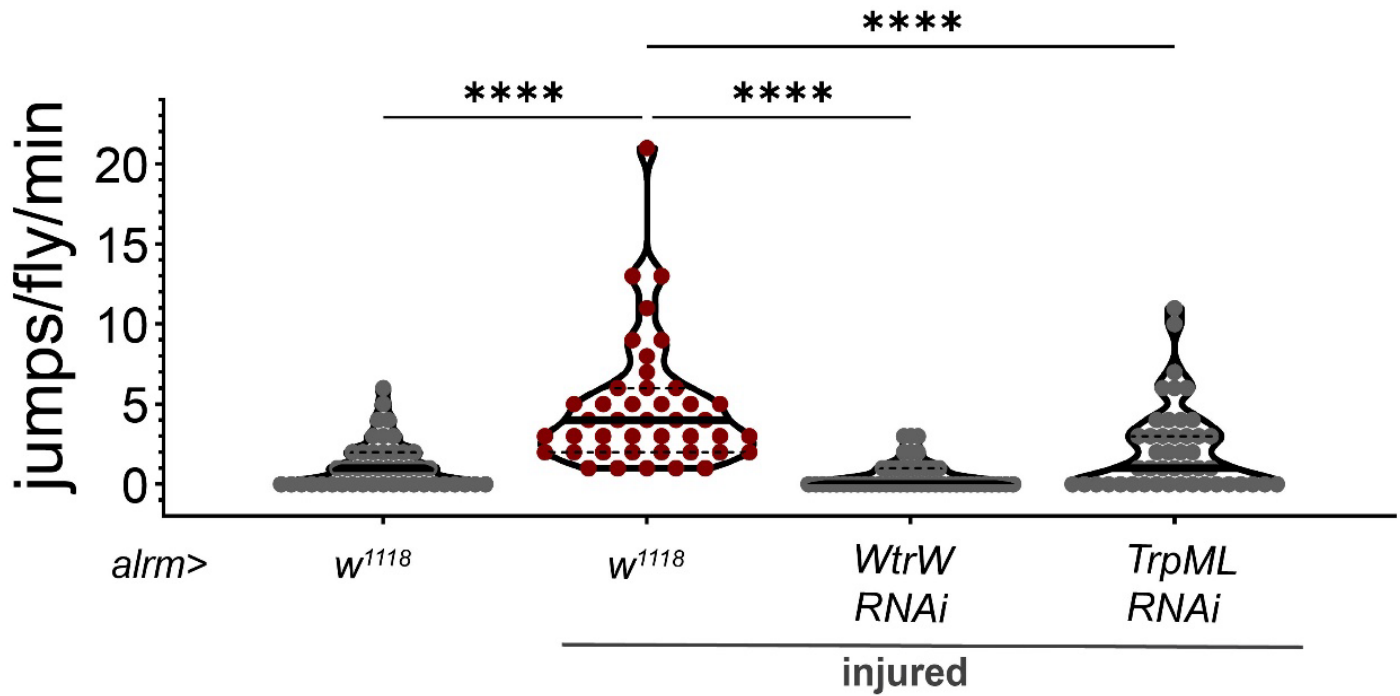

**Supplemental Figure S7. Suppression of astrocyte Waterwitch and TrpML channels attenuates thermal hypersensitivity following injury.** Plot of the number of jumps per fly per minute at 38° C for seven days following leg amputation injury (“injured”) for the indicated genotypes. Each symbol represents the total number of jumps for a single animal ($n=43$ for *alrm-GAL4*; *w<sup>1118</sup>*, $n=45$ for *alrm>Wtrw RNAi*, $n=46$ for *alrm>TrpML RNAi* injured groups and $n=45$ for *alrm-GAL4*; *w<sup>1118</sup>* sham controls) \*\*\*\*, $P<0.0001$; Kruskal-Wallis with Dunn’s multiple comparisons, $N=179$, $H(3)=64.62$, $P<0.0001$.

**Supplemental Table 1. STAR Methods**

| Reagent or Resource | Source | Identifier |
| --- | --- | --- |
| ***Drosophila* Stocks** |  |  |
| w[*]; P{w[+m*]=alm-GAL4.D}3/CyO; Dr[1]/TM3, Sb[1] | BDSC<br>RRID:SCR_006457<br>https://bdsc.indiana.edu/ | astrocyte-like glia<br>GAL4 driver<br>stock# 67031 |
| w[*]; wg[Sp-1]/CyO;<br>P{w[+m*]=alm-GAL4.D}2 | BDSC<br>RRID:SCR_006457<br>https://bdsc.indiana.edu/ | astrocyte-like glia<br>GAL4 driver<br>stock# 67032 |
| w[*]; P{w[+mC]=ppk-GAL4.G}2 | BDSC<br>RRID:SCR_006457<br>https://bdsc.indiana.edu/ | pickpocket sensory<br>neuron GAL4 driver<br>stock# 32078 |
| y[1] v[1]; P{y[+t7.7]<br>v[+t1.8]=TRiP.JF02461}attP2 | BDSC<br>RRID:SCR_006457<br>https://bdsc.indiana.edu/ | UAS TrpA1 RNAi<br>stock# 36780 |
| y[1] sc[*] v[1] sev[21];<br>P{y[+t7.7]<br>v[+t1.8]=TRiP.HMC03562}attP4<br>0 | BDSC<br>RRID:SCR_006457<br>https://bdsc.indiana.edu/ | UAS Orai RNAi<br>stock# 53333 |
| UAS-dOrai(G170M)/cyo, GFP | internal | UAS dOrai G170M |
| UAS-dOrai+/TM6, Tb | internal | UAS dOrai+ |
| y[1] v[1]; P{y[+t7.7]<br>v[+t1.8]=TRiP.JF01957}attP2 | BDSC<br>RRID:SCR_006457<br>https://bdsc.indiana.edu/ | UAS Itpr RNAi<br>stock# 25937 |
| y[1] v[1]; P{y[+t7.7]<br>v[+t1.8]=TRiP.JF02567}attP2 | BDSC<br>RRID:SCR_006457<br>https://bdsc.indiana.edu/ | UAS Stim RNAi<br>stock# 27263 |
| w[*] P{y[+] w[+]=10XUAS-IVS-<br>mCD8::RFP}attP18 P{y[+]<br>w[+]=13XLexAop2-<br>mCD8::GFP}su(Hw)attP8;<br>P{y[+] w[+]=UAS-<br>MKII::nlsLexADBDo}attP40,<br>P{y[+] w[+]=UAS-<br>p65AD::CaM}attP24/CyO;<br>M{w[+]=UAS-p65AD::CaM}ZH-<br>86Fb | BDSC<br>RRID:SCR_006457<br>https://bdsc.indiana.edu/ | TRIC<br>LexAop mCD8-tagged GFP, UAS<br>mCD8-tagged RFP;<br>UAS<br>lexA(DBD)::CaMTP,<br>UAS p65(AD)::CaM;<br>UAS p65(AD)::CaM<br>stock# 62827 |
| y[1] sc[*] v[1] sev[21];<br>P{y[+t7.7] v[+t1.8]=VALIUM20-<br>GAL4.2}attP2 | BDSC<br>RRID:SCR_006457<br>https://bdsc.indiana.edu/ | UAS GAL4 RNAi<br>stock# 35783 |
| **Immunohistochemistry** |  |  |
| Mouse Bruchpilot nc82-s | DSHB | Cat# nc82, 4/16/20<br>RRID:AB_2314866 |
| Rabbit Living colors dsRed | Takara Bio | Cat#632496, Lot<br>2103116 |
|  |  | RRID:AB_10013483 |
| Rabbit anti-GABA | Sigma | Cat#0000154656 |
| Chicken anti-GFP | Abcam | Cat#AB13970 |
| Alexa fluor 488 goat anti-chicken | Invitrogen | Cat#A11039, Lot 1637891 |
| Alexa fluor 568 goat anti-rabbit | Invitrogen | Cat#A11011, Lot 2379475 |
| Alexa fluor 488 goat anti-rabbit | Invitrogen | Cat#A11008, Lot 2420731 |
| Alexa fluor 568 goat anti-mouse | Invitrogen | Cat#A11004, Lot 2332536 |
| **Mounting Reagents** |  |  |
| Kodak Photo-Flo 200 Solution | Electron Microscopy Sciences | Cat#74257 |
| Poly-L-lysine hydrobromide (PLL) | Sigma | Cat#P1524-25MG |
| DPX mountant for microscopy | Electron Microscopy Sciences | Cat# 13512 |
| Ethanol >99.5% (200 proof) | Sigma | Cat# 459844-1L |
| Paraformaldehyde (PFA) 16% Aqueous Solution | Electron Microscopy Sciences | Cat#15710 |
| Phosphate buffered saline (PBS) 1X | Cellgro | Cat#21-040 |
| Triton X-100 | Sigma | Cat#X100 |
| Sylgard 184 silicone elastomer kit | Sigma | Cat# 761028-5EA |
| Schneider's Insect Medium (S2) | Fisher Scientific | Cat#21720-024 |
| Fetal Bovine Serum |  | Cat# |
| **Software** |  |  |
| MATLAB (R2019B) | www.mathworks.com/matlab | RRID:SCR_001622 |
| Fiji | http://fiji.sc | RRID:SCR_002285 |
| R (4.2.2) | https://cran.r-project.org/ | RRID:SCR_001905 |
| CepLDA (R package) | (Krafty, RT, 2016) |  |
| GraphPad Prism (9.5.0 (730)) | www.graphpad.com | RRID:SCR_002798 |
| **Deposited data** |  |  |
| Raw and analyzed data | Figshare |  |
| Analyzed data and codes | GitHub |  |
| **Equipment** |  |  |
| Nikon A1R Confocal Laser Microscope | www.microscope.healthcare.nikon.com | RRID:SCR_020318 |
| Thermal testing 3 mm ring | GitHub |  |
| Vellum translucent paper | Staples | Cat#496755 |
| Isoblock Dry Bath Incubator | Benchmark Scientific | SKU#BSH6000 |
| Fisherbrand Traceable Infrared Thermometer Gun | Fisher Scientific |  |
| Wide 28 mm 77° f/1.7 lens video camera with Sony Exmor IMX363 12.2-megapixel sensor | https://store.google.com | Pixel 4 |
| Dumont #5 forceps | Fine Science Tools | Cat#11252-20 |
| Corning cover glass No.1 | Fisher Scientific | Cat#12-553-451 |
